# *Brucella abortus* egresses from host cells in infective clumps through an actin-dependent mechanism

**DOI:** 10.1101/2025.10.15.682733

**Authors:** Jose Fabio Campos-Godínez, Alejandro Hernández-Saborío, Tiffany Vargas-Moya, Valeria Gómez-Vargas, David Espinoza-Villagra, Ignacio Sandoval, María Paula Rojas-Salas, Melany López-Hernández, Rodrigo Vega-Arce, Reynaldo Pereira-Reyes, Monica Prado, Carlos Chacón-Díaz, Edgardo Moreno, Esteban Chaves-Olarte, Pamela Altamirano-Silva

## Abstract

*Brucella abortus* is an intracellular pathogen whose cell cycle encompasses attachment, internalization, trafficking, replication, and egress from cells. While the intracellular life of *Brucella* has been extensively studied, the mechanisms underlying its exit from host cells are still unclear. In this work, we observed a significant increase in the formation of *B. abortus* containing vacuoles intracellular clumps (BCVIC) and abundant *B. abortus* containing vacuoles-derived extracellular clumps (BCVEC) after 72 h of infection. Membrane extensions protruding through the polymerization of actin filaments were evident in cells at later stages of infection. Purified BCVIC and BCVEC were of similar sizes and predominantly acidic. These vacuoles exhibited a compact arrangement of well-ordered bacteria, comprised of a heterogeneous population of dead and live bacteria and host components LAMP-1 and actin filaments. A proportion of BCVEC were naked, while others were enclosed within an impermeable host membrane. The actin cytoskeleton was implicated in the protrusion of BCVEC since the modulation of Rho GTPases affected the egress of BCVEC. The BCVEC protruding from cells were capable of invading non-infected cells, initiating a new cycle of infection. The topological structure and function of BCVEC underscore their significance in the *Brucella* life cycle as vehicles for bacterial dissemination to host cells and organs.

**Importance:** *B. abortus* is an intracellular pathogen that traffics to the endoplasmic reticulum, where it replicates. However, the mechanisms by which *Brucella* exits host cells and infects new ones are not fully understood. We found that protruding acidic autophagic-like vesicles containing compact, well-organized *Brucella* clusters are shed from cells via GTPase and actin filament recruitment. Through this process, the vesicles are impermeable to antibodies and other substances, like antibiotics, and are highly infectious to neighboring cells. This previously unreported mechanism enhances understanding of the well-known *Brucella* stealth strategy to evade the immune system and establish harmful, long-lasting infections.

## Introduction

*Brucella abortus* is a Gram-negative, intracellular, facultative-extracellular bacterium that causes a long-lasting brucellosis disease (1). In cattle, *B. abortus* induces abortions, low milk production, orchitis, epididymitis, and infertility. In humans, *B. abortus* transmission occurs through direct contact with infected animals or their contaminated products, resulting in a debilitating disease that can be fatal if untreated (2).

Antibody or complement opsonized *B. abortus* can be internalized through zipper phagocytosis by macrophages and monocytes. Alternatively, non-opsonized *Brucella* can enter via fibronectin and lectin receptors or through alternative mechanisms that are not yet fully elucidated (3). *Brucella* internalizes epithelial cells by recruiting actin filaments and small GTPases such as Cdc42, Rac, and Rho (3,4). Inside phagocytic or nonphagocytic cells, *Brucella* is contained within a membrane-enclosed compartment known as the *Brucella*-containing vacuole (BCV) (5,6).

The BCV interacts with early and late endosomes, acidifying the compartment to a pH of ∼4.5 (7); shifting into an early acidified endosomal BCV (BCVe) (8). Then, the BCVe progressively loses endosomal markers while interacting with the exit sites of the endoplasmic reticulum (ER), acquiring markers of this compartment such as calreticulin, calnexin, and sec61β. At this stage, *Brucella* organisms initiate replication (BCVr) (8). Following extensive bacterial replication, the BCVr is captured within acidic autophagosome-like structures, becoming autophagic BCV (8). At later times of infection, *B. abortus* sequesters multivesicular bodies (9), which are generated from the invagination and budding of the limiting membrane (10). Multivesicular bodies can fuse with autophagosomes, forming vesicles called amphisomes that subsequently fuse with the plasma membrane, thereby allowing bacterial egress in multicellular organisms (9). It has been shown that the NIP3L-mediated mitophagy pathway and mitochondrial fragmentation are required to complete the bacterial intracellular cycle and bacterial egress from HeLa cells (11). Likewise, in both HeLa cells and macrophages, intracellular *Brucella* organisms appear to manipulate host iron metabolism to modulate bacterial replication and egress from cells (11,12).

Cells heavily infected with *Brucella* can divide without clear signs of cytotoxic effects (13), and some cells, like trophoblast, may show large acidic vacuoles with clumps of bacteria displaying LAMP-1 markers (14). Here, we demonstrate that, similar to BCV intracellular clumps (BCVIC), the purified BCV extracellular clumps (BCVEC) released from infected cells are characterized by low pH, LAMP-1 and actin filament markers. While some BCVEC protruding from cells are surrounded by a membrane, others are “naked” and released from host cells through the concourse of actin filaments and small GTPases of the Rho subfamily. The released BCVEC successfully infect new host cells, promoting the dispersion of *Brucella* and likely evading the immune system.

## Results

### The exit of *B. abortus* from host cells over time is dynamic

HeLa cells were infected with *B. abortus* 2308W, and the number of intracellular and extracellular bacteria was determined over time (Fig. 1). The exit of *B. abortus* is a dynamic process increased by bacterial replication, with its maximum estimated time reached at 72 hours p.i.

**Figure 1.**
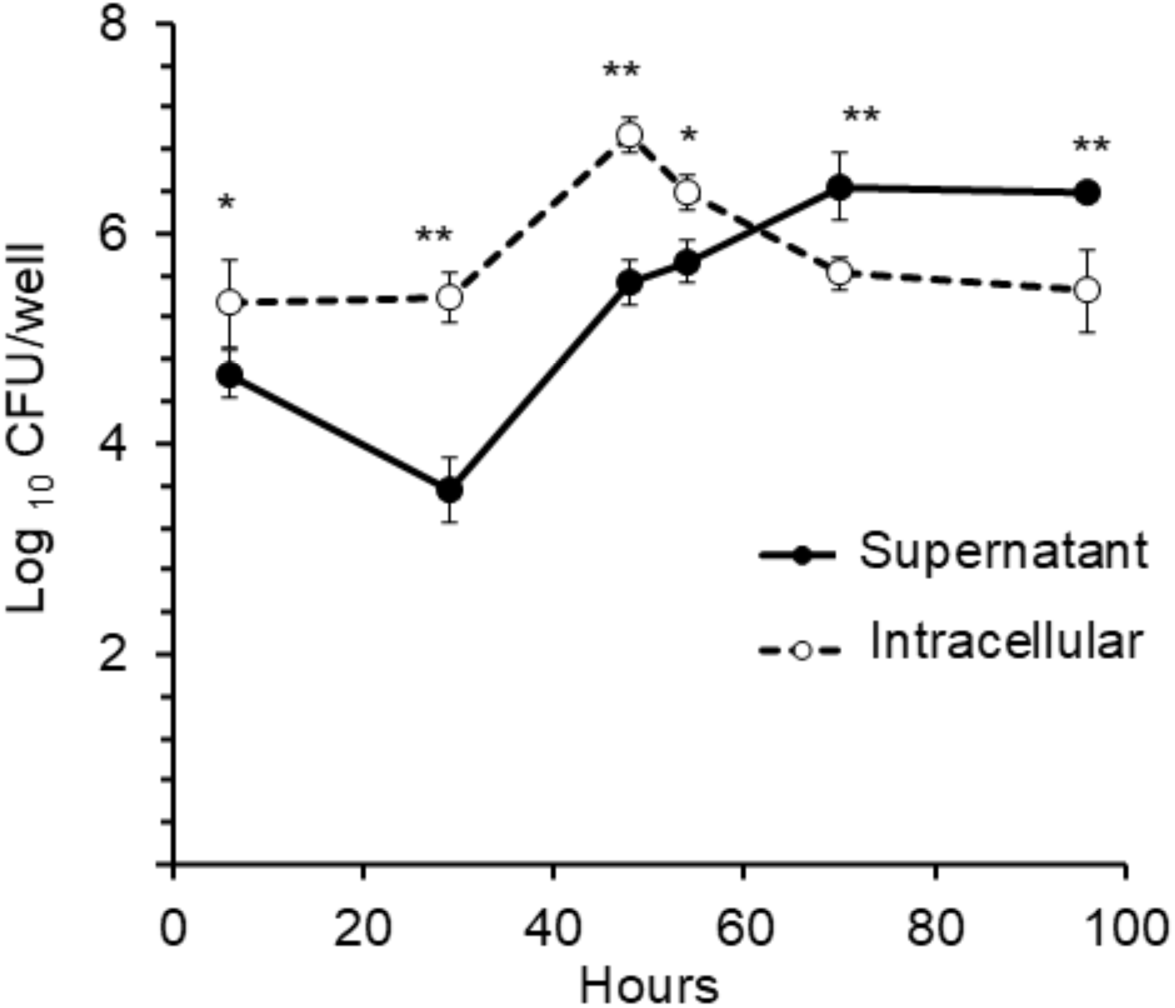
*B. abortus* exit from host cells is a dynamic process. HeLa cells were infected with an early-exponential phase of *B. abortus* 2308W. At the indicated times, the infected cells were washed then incubated for 4 hours in DMEM without gentamicin. Bacteria from the supernatant were collected, plated and CFU counted. Then, cells were washed with DMEM and lysed with 0.1% Triton, and the intracellular CFU determined. Each value is the average of at least three independent determinations.

### *B. abortus* arranges in two different replicative patterns within cells

HeLa cells were infected with *B. abortus*-RFP, and bacteria and infected cells were visualized by fluorescence microscopy over time. No significant differences in the proportion of HeLa cells replicating *B. abortus* were observed between 24 and 72 hours p.i. (Fig. 2A). However, two distinct replicative patterns were noted: i) infected cells with bacteria homogeneously dispersed throughout the cytoplasm (Fig. 2B) and ii) cells with intracellular bacterial clumps named BCVIC (Fig. 2C). From 24 to 66 hours p.i., 70-100% of the cells containing replicating bacteria exhibited dispersed bacteria (Fig. 2D). Conversely, a significant increase of BCVIC was observed in cells after 72 hours p.i. Approximately 56% of the infected cells with replicating bacteria exhibited BCVIC at 72 hours p.i. (Fig. 2C, 2D).

**Figure 2.**
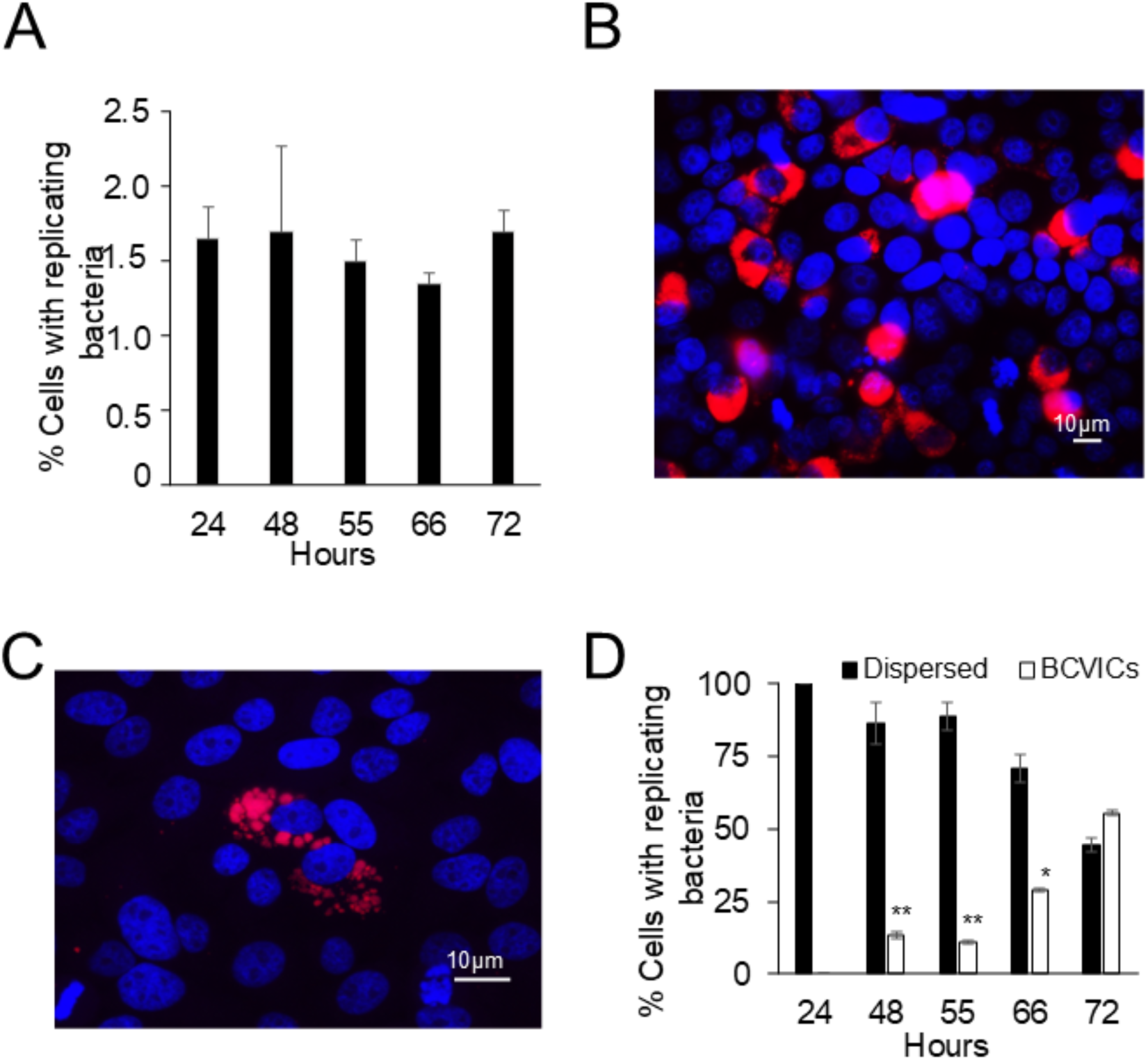
*B. abortus within* intracellular and extracellular vesicles. HeLa cells were infected with an exponential-phase culture of *B. abortus-*RFP (MOI 500). At the indicated times, cells were fixed with paraformaldehyde, and their nuclei were stained DAPI (blue) and visualized by fluorescence microscopy. (A) Proportion of cells with replicating bacteria (>10 bacteria/cell). (B). Immunofluorescence of cells with replicating bacteria dispersed throughout the cytoplasm. (C) Immunofluorescence of cells with BCVIC. (D) Proportion of cells with dispersed bacteria and BCVIC. Each value is the average of at least three independent determinations.

### *B. abortus* organisms protrude from infected cells at later stages of infection

It was observed that at later times of infection, *B. abortus* protrudes from infected cells as single organisms or as BCVEC of different sizes (Fig. 3A, 3B, 3C). The protruding BCVECs of bacteria may be “naked” or surrounded by a host membrane structure (Fig. 3C). Scanning electron microscopy revealed that bacteria in BCVEC arranged in closely packed, organized clumps (Fig. 3D) interlinked by thin structures (Fig. 3E). The proportion of infected cells with BCVECs protruding after 80-83 hours p.i was close to 20%, and egress was not significantly affected by the presence of gentamicin in the culture media, suggesting that some of the BCVEC originating from BCVIC had a protective membrane derived from the host cell (Fig. 3F), a phenomenon confirmed by electron microscopy (Fig 3G an H

**Figure 3.**
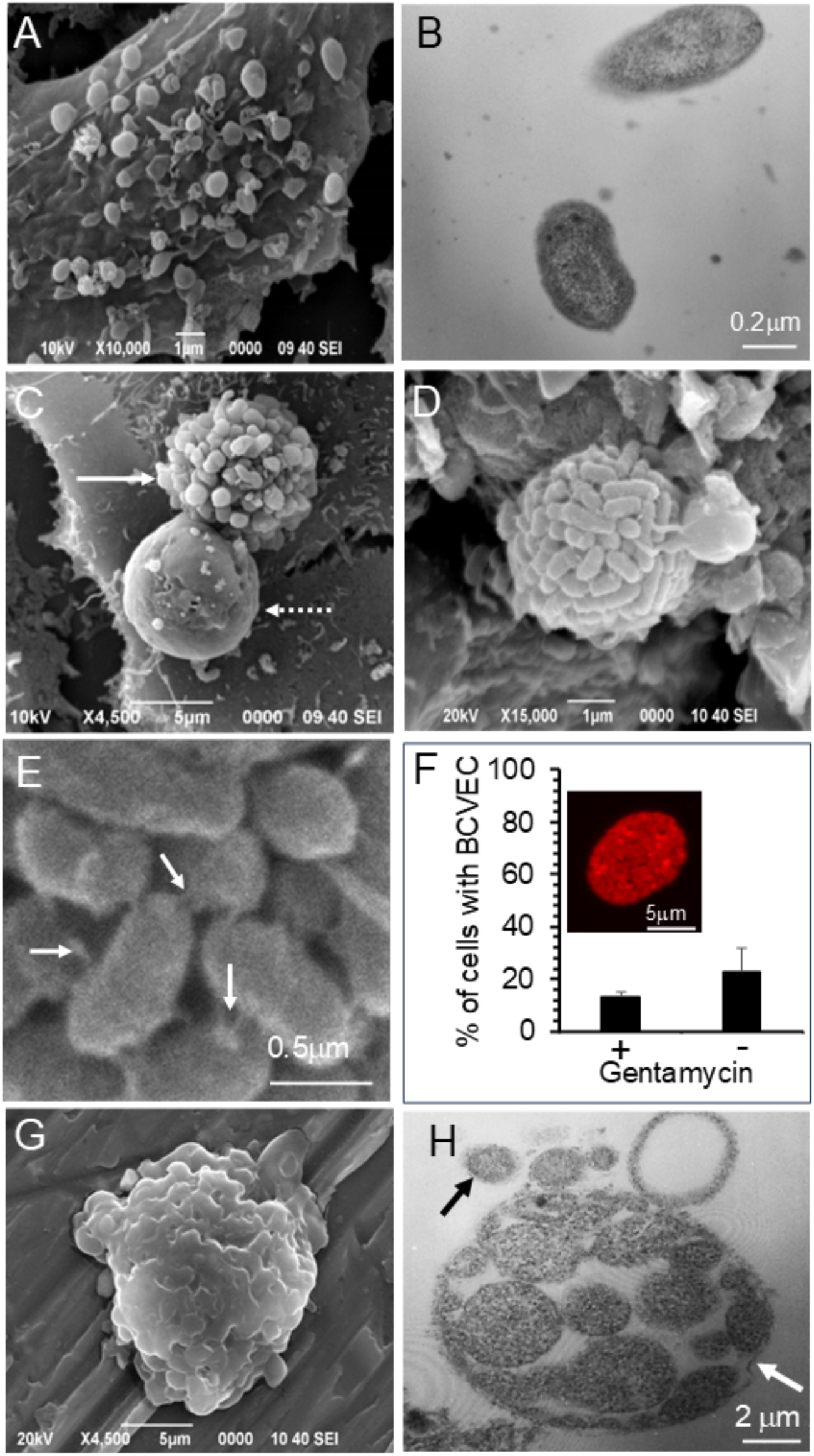
*B. abortus* protrude from cells as single bacteria, naked BCVEC or membrane-enclosed BCVEC. HeLa cells were infected with an exponential-phase culture of *B. abortus*-RFP (MOI 500). After 72 h, cells were incubated without antibiotics for 4 hours, fixed and processed for scanning electron microscopy or fluorescence microscopy. (A) Scanning electron microscopy of individual bacteria protruding from infected HeLa cells. (B) Transmission electron microscopy of single bacteria released by infected HeLa cells. (C) Scanning electron microscope of two forms of BCVEC protrusion: “naked” BCVEC (continuous white arrow) or membrane-enclosed BCVEC (discontinuous with arrow). (D) Protruding naked BCVEC are arranged in closely tightened aggregates of bacteria. (E) Interbacterial linking structures within the bacterial aggregates (white arrows). (F) From 48-100 hours p.i., infected cells were monitored by live imaging, and the proportion of BCVEC protruding from infected cells was determined in the presence or absence of gentamycin. The insert (in red) shows a fluorescent BCVEC filled with *B. abortus*-RFP secreted by an infected cell after 100 hours of infection. (G) Scanning electron microscopy of released BCVEC.(H) Transmission electron microscopy of purified BCVIC. Notice the thin cell-derived membrane surrounding the BCVIC (white arrow) and a free bacterium (black arrow).

### A host-derived membrane surrounds a significant proportion of BCVEC and BCVIC

Analysis of the supernatants of *B. abortus*-RFP HeLa-infected cells shows the release of individual bacteria and BCVEC. In each 60X field examined, ∼1000 free bacteria and at least one BCVEC were present. To determine the proportion of BCVEC covered by a host-derived membrane, purified vesicles were incubated with anti-*Brucella* LPS-FITC antibody under non-permeabilizing conditions. Approximately 70% of the BCVEC were stained with anti-*Brucella*-LPS antibodies, indicating that the remaining ∼30% were surrounded by a host impermeable membrane (Fig 4A and B). In addition, a significant proportion of purified BCVIC was also covered by a host-derived membrane (Fig 4A). These findings were corroborated by labeling the BCVEC with wheat germ agglutinin conjugated with Alexa Fluor 488, which primarily reacts with N-acetyl-D-glucosamine and sialic acid residues of glycoproteins on cell membranes (Fig. 4C). Overall, the results indicate that the BCVEC are a heterogeneous population of vesicles, with one-third of them surrounded by an antibody-impermeable membrane. Those BCVEC permeable to antibodies may still be partially surrounded or contain disrupted membranes.

**Figure 4.**
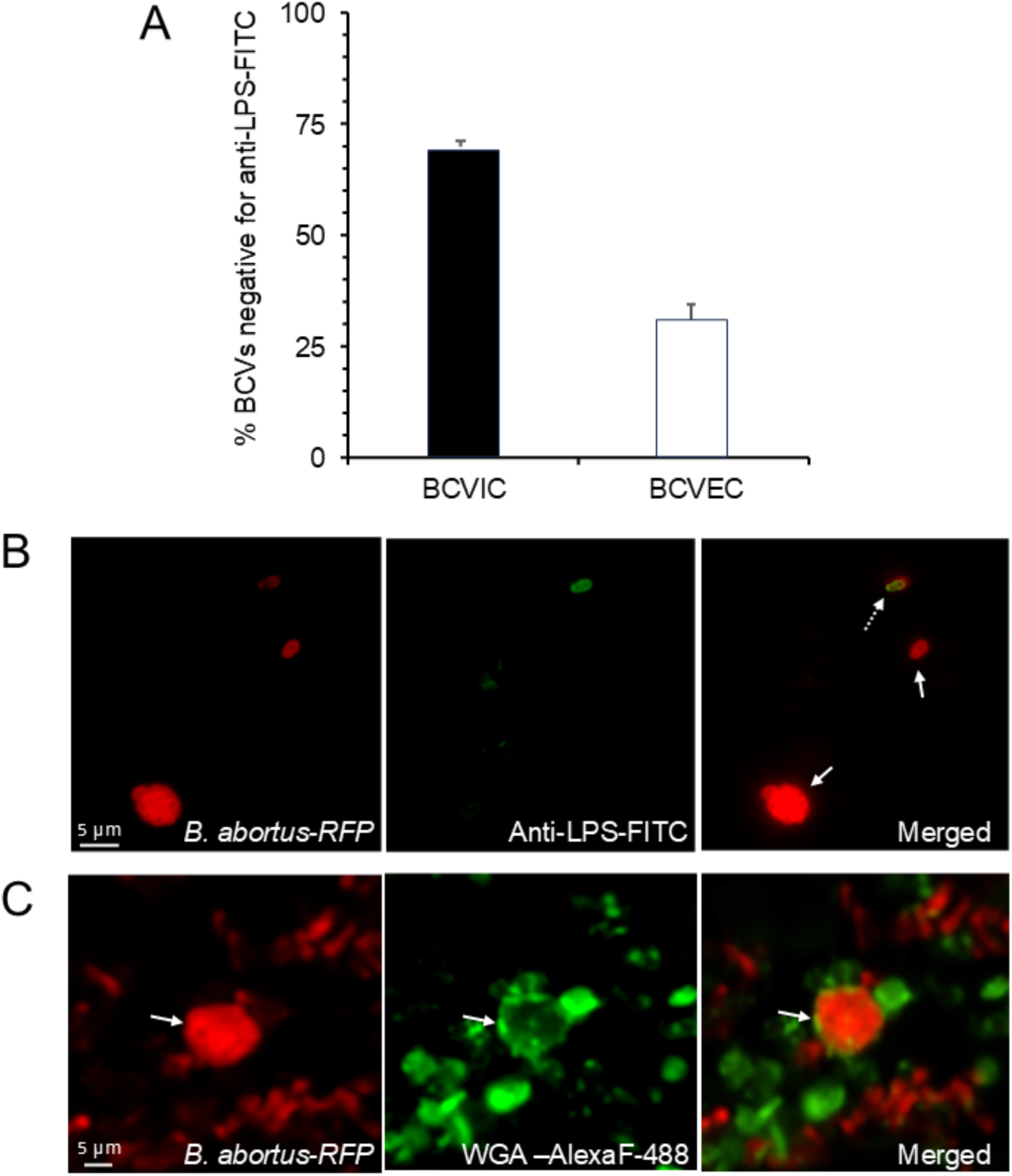
The membrane-surrounded BCVEC and BCVIC are impermeable to antibodies. HeLa cells were infected with an exponential-phase culture of *B. abortus*-RFP (MOI 500). After 72 hours of incubation, cells were incubated without antibiotics for 4 hours, and the supernatant was collected. The cells were then lysed, and BCVIC and BCVEC were purified by centrifugation cycles and stained with fluorescent conjugates. (A) Purified BCVIC and BCVEC were stained with anti-*Brucella* LPS-FITC without permeabilization and counted. Each value is the average of at least three independent determinations. (B) Purified BCVIC stained with anti-*B. abortus*-LPS-FITC. The discontinuous arrow points to a bacterium not surrounded by an impermeable host cell membrane, while the continuous arrow points to a BCVIC and a bacteria impermeable to anti-*B. abortus*-LPS-FITC. (C) Purified BCVEC were stained with wheat germ agglutinin-Alexa Flour-488 (WGA-alexaF-488, green). White continuous arrows indicate the BCVEC surrounded by an impermeable membrane. The contrast was adjusted to no more than 20% in the same proportion in the respective images using the Hue adjustment function of Adobe Photoshop ®. Raw images were processed using the Denoise.ai module.

### BCVEC and BCVIC size and shape

The average volume of an uninfected HeLa cell ranges from 2500-3000 μm^3^, a volume which may increase in infected cells filled with non-toxic *Brucella* organisms (13,15). Consequently, infected cells containing a significant number of BCVIC may release BCVEC of various sizes, containing just a few or hundreds of bacteria. The analysis of the BCVEC and BCVIC observed by immunofluorescence revealed that these vesicles may accommodate from ∼10 to ∼650 bacteria (Fig. 5A). The more abundant BCVEC and BCVIC carry less than 300 bacteria. Vacuoles above 250 μm^3^ are significantly less abundant than smaller ones and probably more prone to disruption due to lower pressure and a larger surface area-to-volume ratio.

**Figure 5.**
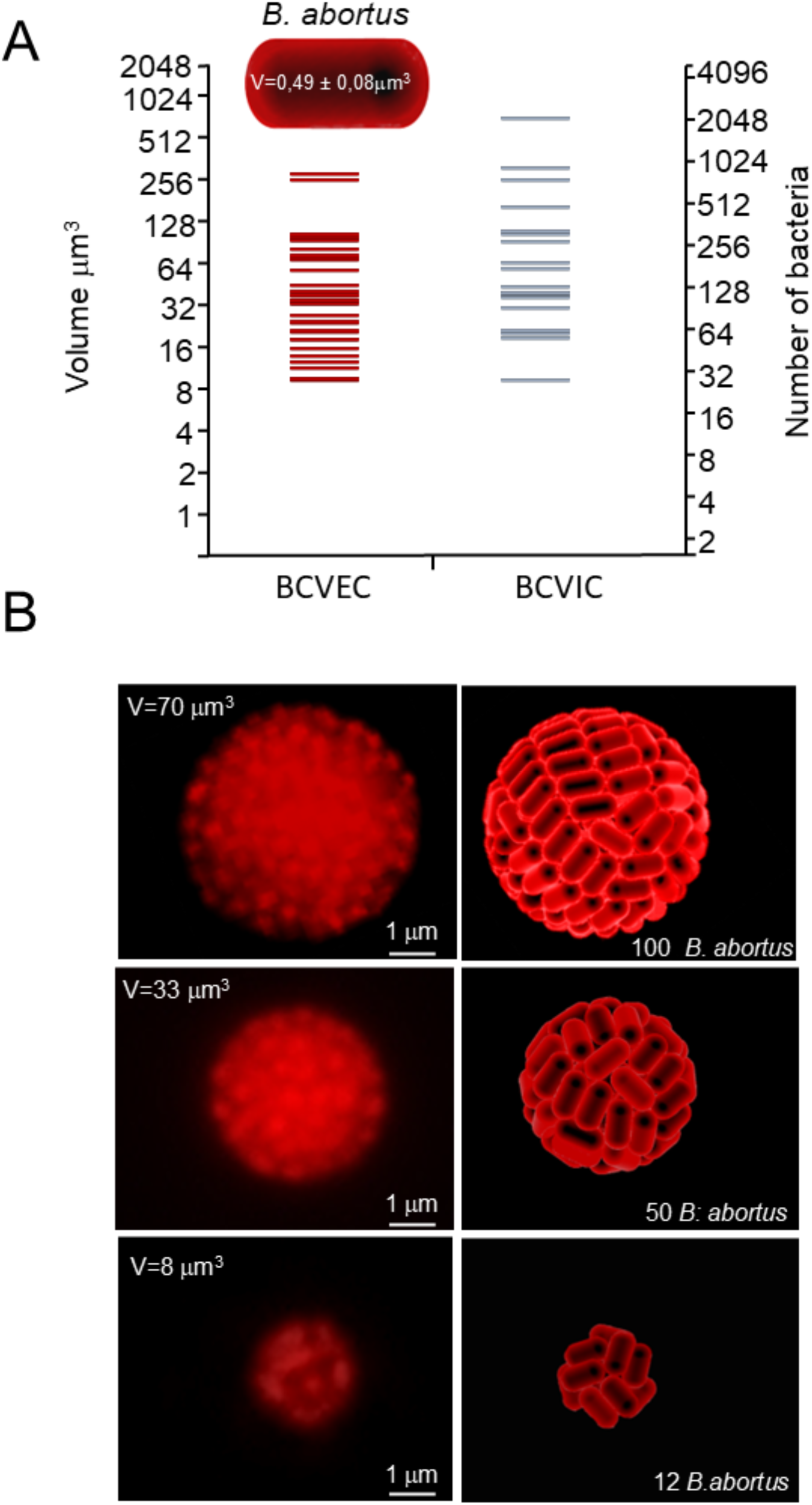
Structural arrangement of BCVEC and BCVIC. Purified BCVIC and BCVEC were counted and observed under a confocal fluorescence microscope. (A) Volumes of BCVEC and BCVIC, and number of *B. abortus*-RFP per vacuole. The *B. abortus* volume estimation was 0.49±0.08 μm^3^ (inserted model). (B) BCVEC of different sizes and the corresponding packing and number of *B. abortus*-RFF per BCVEC of a given volume were estimated. The photographs of the vacuoles were amplified (left panels), the number of individual *B. abortus*-RFP calculated using the Adobe Photoshop ® density color pixel function of the image’s sections, and the packing of bacteria (l/d=1) within vacuoles modeled (right panels) following the method for close packing of short rods within spherical surfaces (16). The contrast was adjusted to no more than 20% in the same proportion in the respective images using the Hue adjustment function of Adobe Photoshop.

Amplification of BCVEC photographs of various sizes revealed a regular arrangement of bacteria within the vacuoles (Fig. 5B). This action allowed their analysis by color density differences through the topology of the BCVEC photographs and the estimation of the approximate number of bacteria within the vacuoles. Following the estimation method of the close packing of short rods within spherical surfaces (16), we modeled the packing of *B. abortus* coccobacilli within BCVEC of three different sizes (Fig. 5B). The arrangement disposition of coccobacilli in each of the three BCVEC models considerably differed. For the structures with 50 and 100 coccobacilli, the spherical arrangements were consistent with the BCVEC photographs, as predicted by the theory for packing short rods. However, for the smaller BCVEC, pairs of nearly parallel coccobacilli forming the faces of a cube were observed (Fig. 5B).

### BCVEC and BCVIC contain a heterogeneous population of *B. abortus*

Most bacteria constitutively expressing fluorescent proteins, such as RFP or GFP, maintain cell integrity and are generally considered alive; in contrast to dead bacteria from which fluorescence rapidly decays (17). The fluorescent analysis of BCVEC revealed that they are heterogeneous in terms of bacterial protein metabolism and the living *B. abortus* population (Fig 6A-C). The proportion of detectable *B. abortus*-RFP-pJC45-gfp displaying an inducible GFP (synonymous with metabolically protein synthesis-active bacteria) within BCVEC and BCVIC was not significantly different (Fig. 6D). The fact that close to 50% of the vacuoles do not display detectable GFP does not mean that they are composed of just metabolically inactive bacteria but that the number of active bacteria present within them may be below the threshold of detection and that the expression of the GFP was not fully accomplished by the ATc not readily diffusing inside the vacuoles (Fig 6A). Indeed, the proportion of live bacteria within BCVEC was close to 85%, while the dead were ∼18% (Fig. 6E). Likewise, the analysis of metabolically active and living inactive *B. abortus,* as well as dead bacteria, confirmed that BCVEC is composed of these three bacterial populations in different proportions (Fig. 6F). Altogether, these results indicate that BCVEC contains ∼85% living organisms and 15% dead bacteria; among the living organisms, close to 5% are not metabolically active.

**Figure 6.**
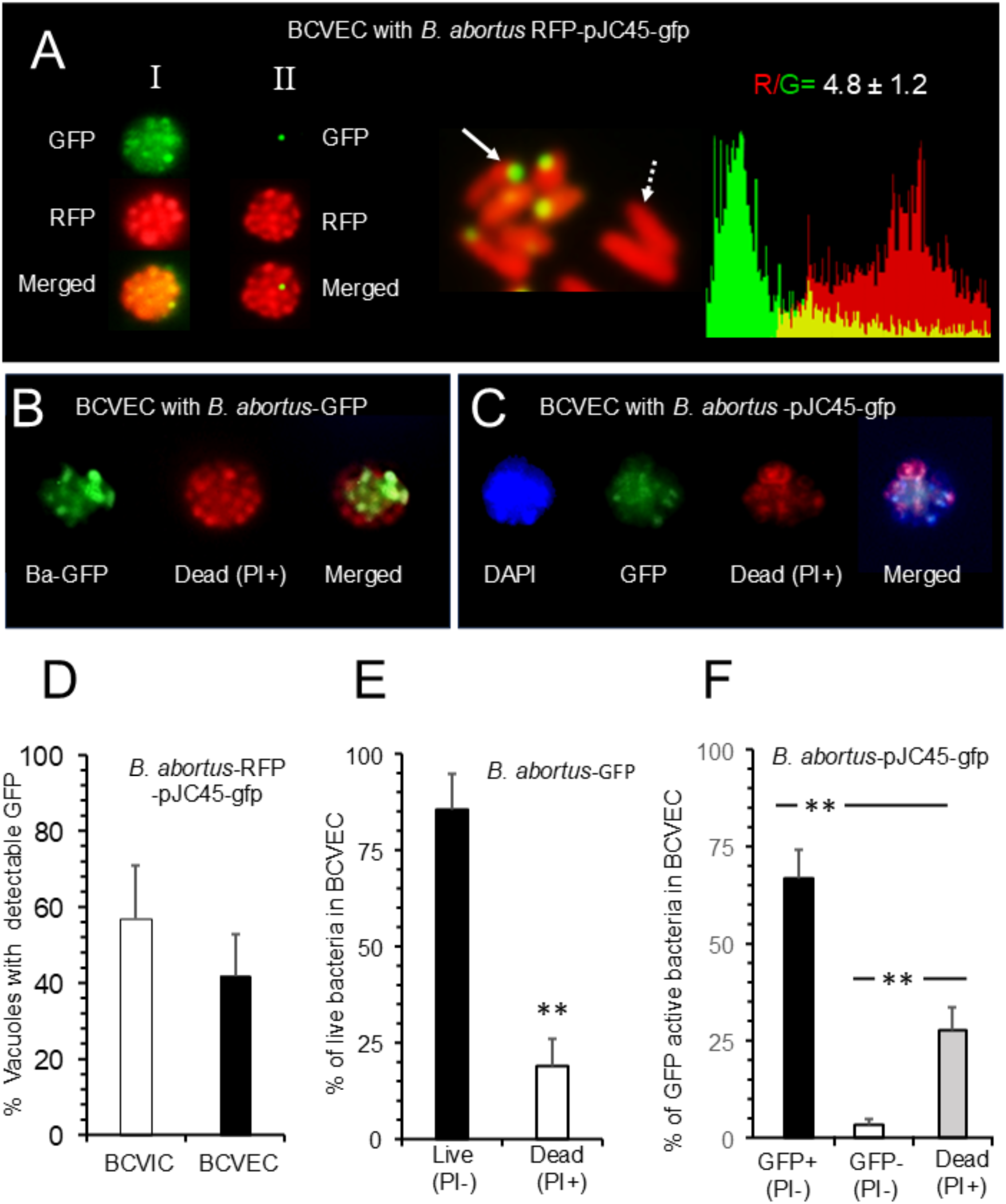
BCVEC and BCVIC have a heterogeneous population of *B. abortus*. HeLa cells were infected with an early-exponential-phase inoculum of either *B. abortus*-GFP*, B. abortus-*pJC45-gfp*, or B. abortus*-RFP-pJC45-gfp (MOI of 500). After 72 hours, the cells were incubated without antibiotics for 4 hours and BCVICs and BCVECs purified and analyzed. (A) To the left of the panel: a BCVEC containing *B. abortus*-RFP-pJC45-gfp with practically all bacteria expressing GFP (I), and a BCVEC containing only one bacterium expressing GFP (II). In the center of the panel: *B. abortus*-RFP-pJC45-gfp expressing GFP (unbroken arrow) and live bacteria non-expressing GFP (broken arrow). To the right a spectrum of one bacterium expressing GFP (unbroken arrow). Based on the analysis of 50 spectra, the ratio between red and green (R/G) fluorescence was estimated. (B) BCVEC containing live *B. abortus*-GFP (Ba-GFP) and dead *B. abortus*-GFP (PI+) organisms. (C) BCVEC with live, metabolic active and dead *B. abortus* (DAPI in blue). In green, the metabolic active *B. abortus*-pJC45-gfp expressing GFP. In red, dead bacteria stained with propidium iodide (PI+). (D) Proportion of BCVEC and BCVIC with at least one metabolically active *B. abortus*-RFP-pJC45-gfp expressing GFP. (E) Purified BCVEC containing *B. abortus*-GFP were lysed with 0.1% Triton, and the bacterial viability was determined using propidium iodide (PI) stain. (F) Purified BCVEC containing *B. abortus-*pJC45-gfp were lysed with 0.1% Triton, and the metabolic activity of bacteria was determined by inducing the production of GFP in *B. abortus*-pJC45-gfp through the action anhydro-tetracycline (250 nM) for six hours. Then, all bacteria were visualized with DAPI, and dead bacteria were detected with propidium iodide (PI). The contrast was adjusted to no more than 20% in the same proportion in the respective images using the Hue adjustment function of Adobe Photoshop ®.

### BCVEC and BCVIC are acidic and harbor LAMP-1 and actin filaments

Consistent with the previously described acidic properties of BCVIC ^8^ and the similar proportion of BCVIC and BCVEC with *B. abortus* expressing GFP (Fig. 6D), we determined that practically all BCVEC were acidic (Fig. 7A). Likewise, all examined BCVIC and BCVEC were labeled with the membrane protein LAMP-1 and phalloidin for actin detection (Fig.7B). Our results indicate that BCVEC originates from BCVIC and that the general properties of the BCVEC are maintained after protrusion from the cell.

**Figure 7.**
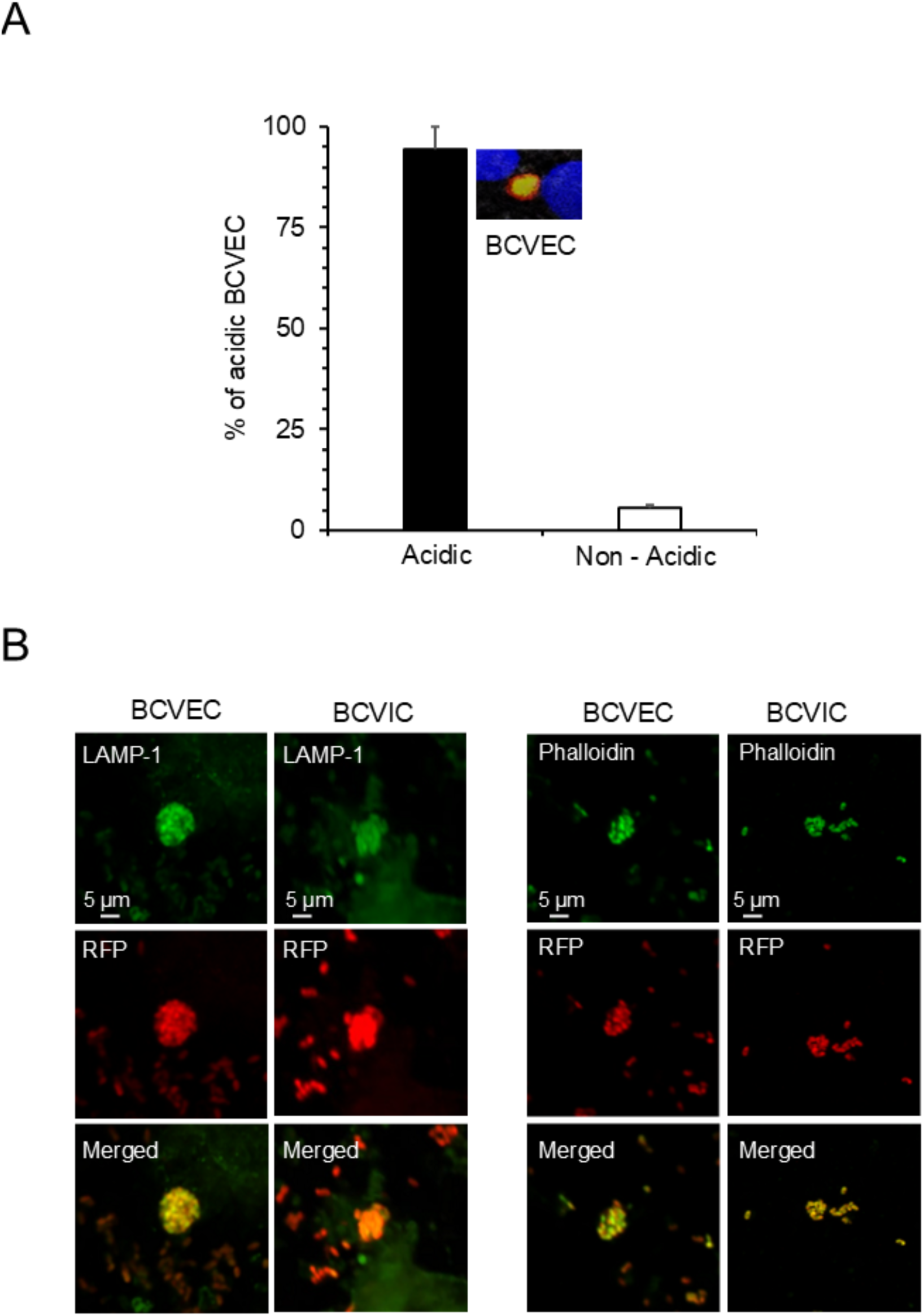
BCVIC and BCVEC are acidic and rich in LAMP-1 and actin filaments. (A) Protruding BCVEC were labeled with 50 nM LysoTracker Red DND-99 (Invitrogen) for 1 hour and observed through the BioTek Lion Heart Imager every 3 hours until 100 hours p.i., and the acidic BCVEC counted. The insert shows a BCVEC in yellow. (B) Purified BCVEC and BCVEC were stained with anti-LAMP-FITC or phalloidin-FITC and observed under the fluorescent microscope. Raw images were processed using the Denoise.ai module.

### BCVEC protrusion is linked to the actin-cytoskeleton polymerization

Single *B. abortus* and BCVEC were observed to emerge from infected cells in association with actin filaments (Fig. 8A and B). As the infection progressed, there was an increase in the proportion of infected cells with actin-mediated cytoplasmic extensions, which are notably shorter and less conspicuous in non-infected cells (Fig. 8C, D, and E). These data indicate a linkage between actin filaments-mediated cell extensions and the protrusion of BCVEC.

**Figure 8.**
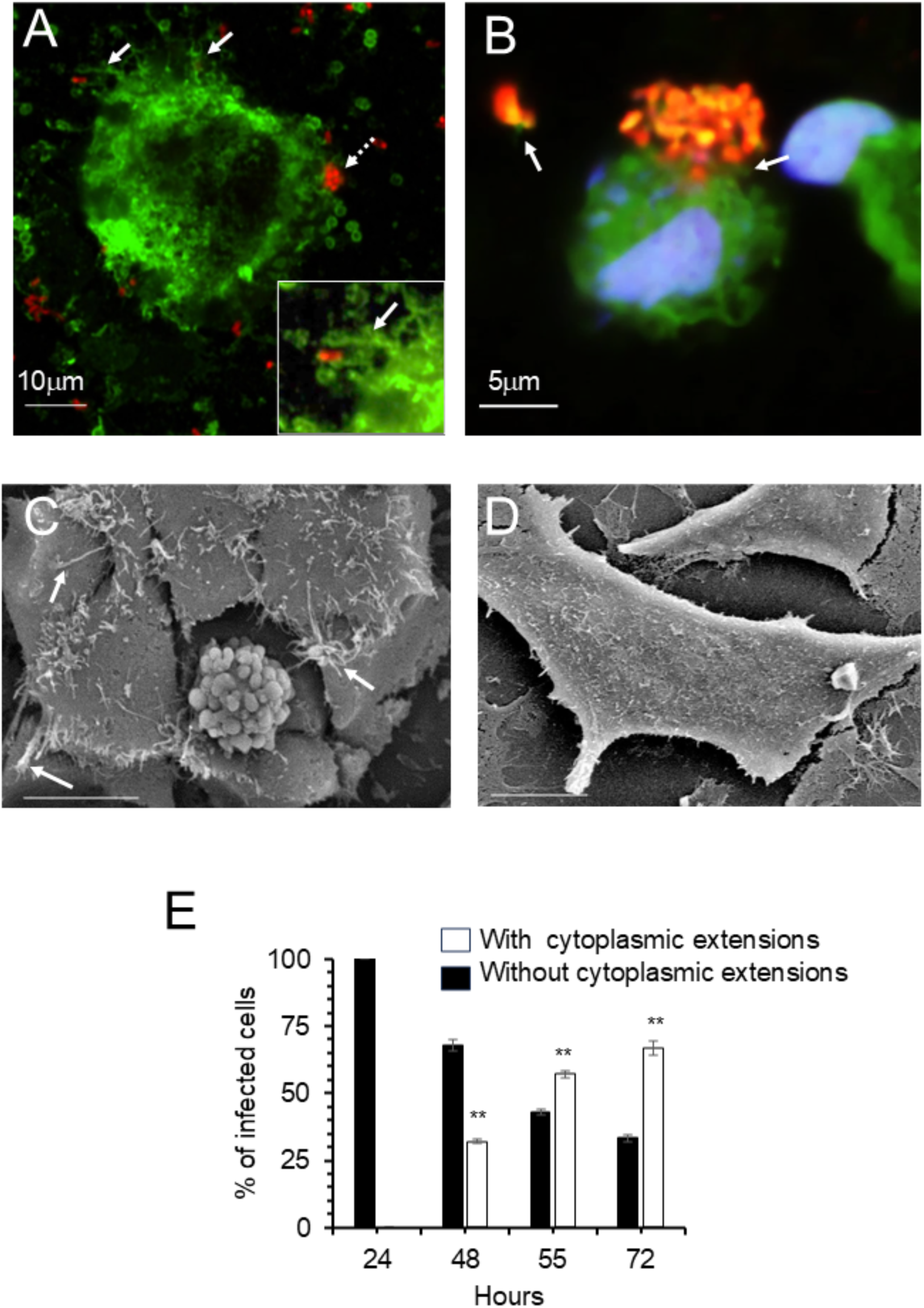
Actin filaments are associated with protruding bacteria and BCVEC. (A) *B. abortus*-RFP protruding from infected HeLa cells (broken arrow) displaying cytoplasmic extensions, stained with phalloidin (unbroken arrows) after 48 hours. The insert is a 2 x amplification of a cytoplasmic insertion.Raw images were processed using the Denoise.ai module. (B) BCVEC (in red) protruding from cells resting over actin cytoskeleton filaments (white arrows) stained with phalloidin (green). The cell nucleus was stained with DAPI (blue). Raw images were processed using the Denoise.ai module. (C) Scanning electron microscopy of infected cells with protruding naked BCVEC and displaying long and numerous thin cytoplasmic extensions (white arrows). (D) Scanning electron microscopy of non-infected HeLa cells. (E) Proportion of infected HeLa cells with cytoplasmic extensions over time after infection. Each value is the average of at least three independent determinations.

### GTPases of the Rho subfamily promote the formation of BCVIC and protrusion of BCVEC

The activation of monomeric GTPases through the addition of CNF significantly enhanced the formation of BCVIC at later times of infection (Fig. 9A). Likewise, CNF also promoted the protrusion of BCVEC from the cells (Fig. 9B). In contrast, inhibition of Rho GTPases with *C. difficile* toxin B, significantly decreases the BCVEC protrusion from the cells (Fig. 9B). These outcomes were consistent regardless of whether the infected cells were incubated with or without gentamicin. Moreover, exposure of infected cells to CNF significantly increases the bacterial release. In contrast, exposure to cytochalasin inhibits bacterial release due to the disruption of the actin cytoskeleton (Fig. 9C). Scanning electron microscopy confirmed that CNF, in addition to promoting the protrusion of BCVEC, also favors the increase of membrane filaments, while Toxin B works in the opposite direction, inhibiting the protrusion of BCVEC and actin-mediated membrane filaments (Fig. 9D). These events demonstrate that recruitment of GTPases and the concomitant actin polymerization are necessary events for the generation of BCVIC and protrusion of bacteria and BCVEC.

**Figure 9.**
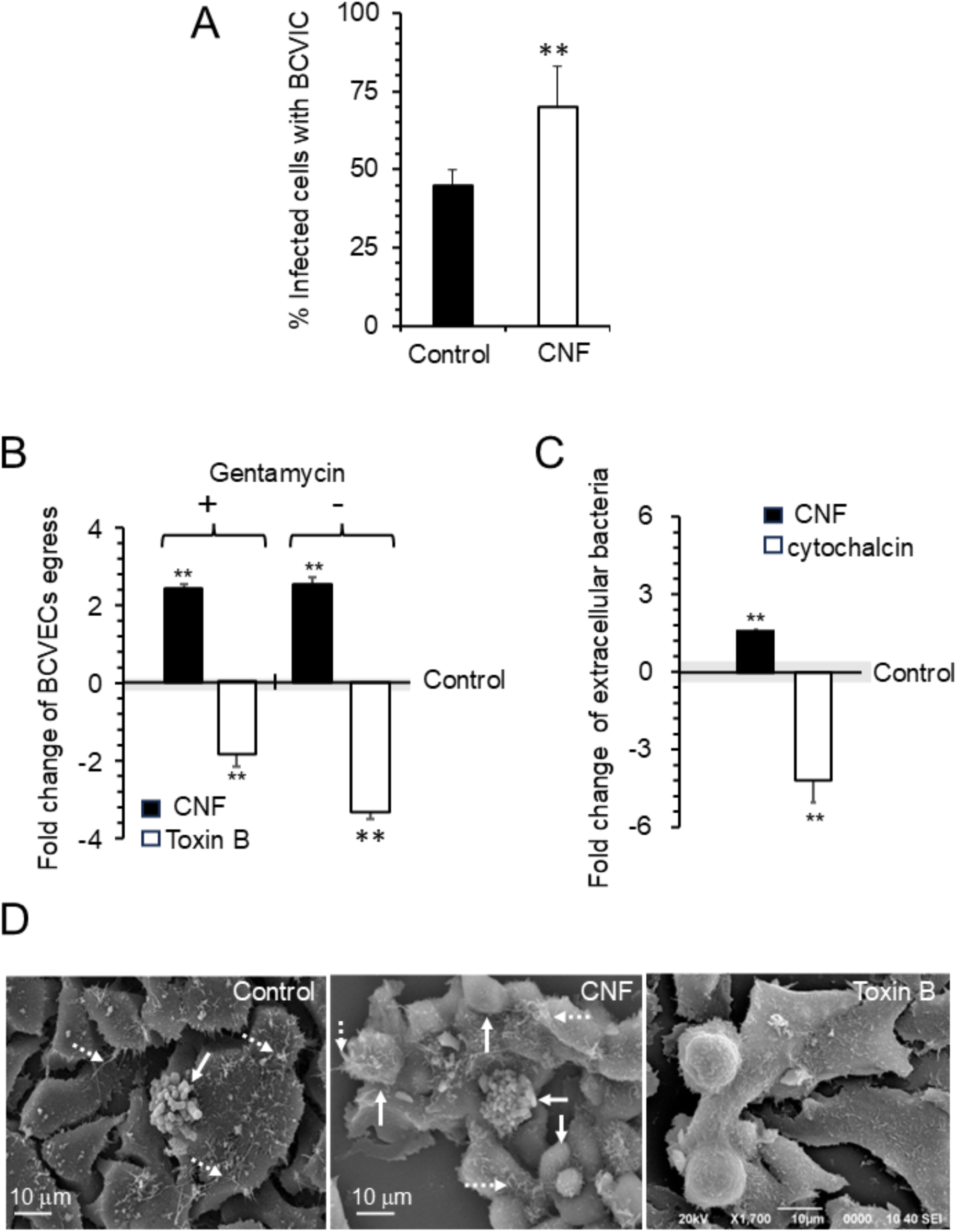
GTPases of the Rho subfamily and actin polymerization promote the exit of BCVEC. HeLa cells were infected with a *B. abortus*-RFP (MOI of 500) and received different treatments after 48 hours post-infection. Then, cells were incubated and data collected after 90 hours of infection. (A) The proportion of *B. abortus*-infected HeLa cells after treatment with CNF. (B) Fold increase and decrease in the protrusion of BCVEC from HeLa cells after treatment with CNF or *C. difficile* toxin B in the presence or absence of gentamycin. The standard error of the control is indicated by the gray area over the “X” axis. (C) Fold increase and decrease of extracellular bacteria concerning the total number of bacteria (intracellular + extracellular) detected in infected HeLa cells treated with CNF or cytochalasin. The standard error of the control is indicated by the gray area over the “0” of the “X” axis. (D) Scanning electron microscopy of a *B. abortus* infected HeLa cell without treatment (Control), treated with CNF, and treated with *C. difficile* toxin B. Notice that the protruding BCVEC (unbroken arrow) is accompanied by numerous long and thin cytoplasmic cell extensions (broken arrows) in the infected HeLa cells, which are more conspicuous in the CNF-treated cells and practically absent in the toxin B-treated infected HeLa cells. Note that the CNF-treated infected HeLa cells show a significant increase in the formation of naked and membrane-bound BCVEC.

### BCVEC contain infective *Brucella*e

To determine whether *B. abortus* organisms inside BCVEC can infect non-infected cells, live imaging was conducted 48 to 100 hours p.i. (Fig. 10A). Remarkably, close to 100 % of the BCVEC initiated a new infection cycle regardless of the presence or absence of gentamicin in the culture media (Fig. 10B). Exposure to propidium iodide revealed that after more than 4 days of culture, the cell membrane integrity remained impermeable in close to 100% of the BCVEC donor-infected cells, in contrast to the non-infected cells. This event indicates that *B. abortus* intracellular proliferation and later BCVEC shedding prolong the life of these infected cells for a protracted period (Fig. 10C and D).

**Figure 10.**
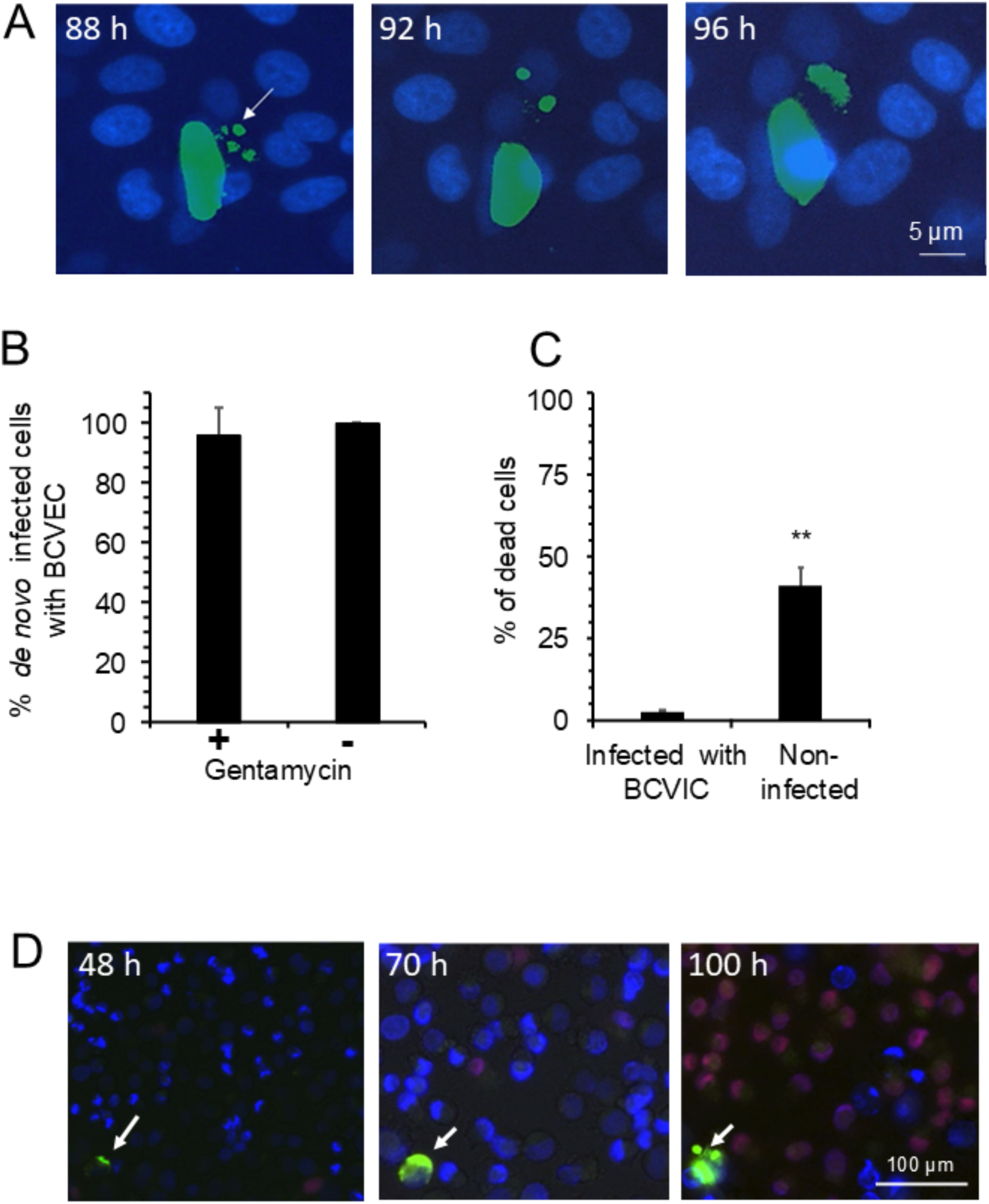
BCVEC shed by HeLa cells initiate a new cycle of infection. (A) BCVEC shedding by an infected HeLa cell (white arrows) and *de novo* infection of naïve cells were monitored using live imaging every 3 hours for 100 hours. *B. abortus*-GFP (in green), cell nuclei (in blue). The frames are the same microscopic field at different times. (B) The proportion of BCVEC that infect new cells and initiate replication was determined in the presence or absence of gentamycin. (C) Proportion of dead cells in non-infected and infected HeLa cells with protruding BCVEC. (D) Fluorescent microscopy of *Brucella*-GFP-infected and non-infected HeLa cells stained with propidium iodide at different times. The *B. abortus*-GFP infected cell (white arrow) remains alive (not stained with propidium iodide) and, in time, shedding BCEVC, while a large proportion of non-infected cells died after 100 hours of culture (stained in red). The frames are the same microscopic field at different times. Each value is the average of at least three independent determinations.

## Discussion

Several strategies for the egress of intracellular bacterial pathogens from the host cells have been reported (5,9,18–21). These pathways include the release of single, naked bacteria following host cell death mediated by toxic bacterial mediators or the cell disruption caused by stress mechanisms due to overcrowding of bacteria within host cells. Alternatively, bacteria may egress by non-lytic mechanisms, either by actively breaching host cell-derived membranes or surrounded by host cell membranes. In this latter mechanism, bacteria may protrude as single cells or as large vesicles filled with bacteria (20). As expected, these mechanisms are not mutually exclusive and can be observed during a cell infection process with a given pathogen (20).

In the case of *Brucella,* it has been shown that heavily *B. abortus*-infected trophoblasts release free bacteria into the uterine lumen after cell necrosis and ulceration of the chorioallantoic membranes, a phenomenon associated with the overwhelming infection of these cells (5,18). Disruption of heavily *Brucella*-infected cells does not result from direct toxic action by the replicating organisms but is likely due to metabolic exhaustion and stress (13,15). Indeed, at later times of infection, *Brucella* organisms occupy most of the cytoplasmic space of the epithelial cells within endoplasmic reticulum-derived structures (5,6,13,18,22). Consequently, one alternative is that *Brucella* organisms trigger an endoplasmic reticulum stress response, which eventually disrupts the heavily infected cell, releasing free bacteria into the milieu (23).

A recent study (24) suggested that during their replication, *Brucella* organisms undergo a conversion to a rough phenotype, inducing cytotoxic factors that trigger cell death and bacterial release. However, this model is unlikely. As stated previously, it has been extensively demonstrated that replicating *Brucella* organisms, in addition to maintaining their smooth phenotype throughout infection (they react with anti-smooth LPS antibodies), and are non-toxic to cells (13,15). Indeed, in agreement with our previous observations, we demonstrate that infected HeLa cells shedding BCVEC survive longer than most non-infected cells in cultures. This event suggests that *Brucella* organisms somewhat arrest the cell death of infected cells for a protracted period, allowing for high intracellular replication and the eventual shedding of highly infectious BCVEC. However, it seems plausible that after a prolonged time (> 100 hours), these heavily infected cells would exhaust and eventually die, releasing bacteria into the surrounding milieu, as previously indicated and observed in infected trophoblasts (5,25).

It has been shown that *Brucella*-containing vacuoles acquire autophagic characteristics (24,26), promoting a non-lytic exocytosis process through the sequestration of multivesicular bodies forming amphisomes that eventually fuse with the plasma membrane and generate bacterial egress as vesicles (9). Here, we have validated this trafficking pattern and extended the findings on the dynamics, composition, and infectious properties of the BCVEC. A model of our proposal for the egress of *Brucella* organisms in BCVEC from infected cells is presented in **Fig. 11**. Following extensive replication in endoplasmic reticulum-derived compartments, *B. abortus* organisms assemble into an acidic vacuole with autophagosomal characteristics (BCVIC), as proposed by Starr et al. (2012). During this period, replicating bacteria prolong the lifespan of the infected cell. The BCVEC formation is initiated by the activation of Rho subfamily GTPases and the active polymerization of actin filaments. The released BCVEC retains the acidic properties and general characteristics of BCVIC. The membrane may protect the bacteria from immune detection and facilitate infection of neighboring cells. Additionally, free *B. abortus* organisms are released into the extracellular milieu. These single bacteria can reach adjacent cells or distant sites, although they are more vulnerable to immune clearance. Infected cells display membrane protrusions, suggesting actin recruitment. This may allow the bacteria to spread while avoiding immune exposure.

**Figure 11.**
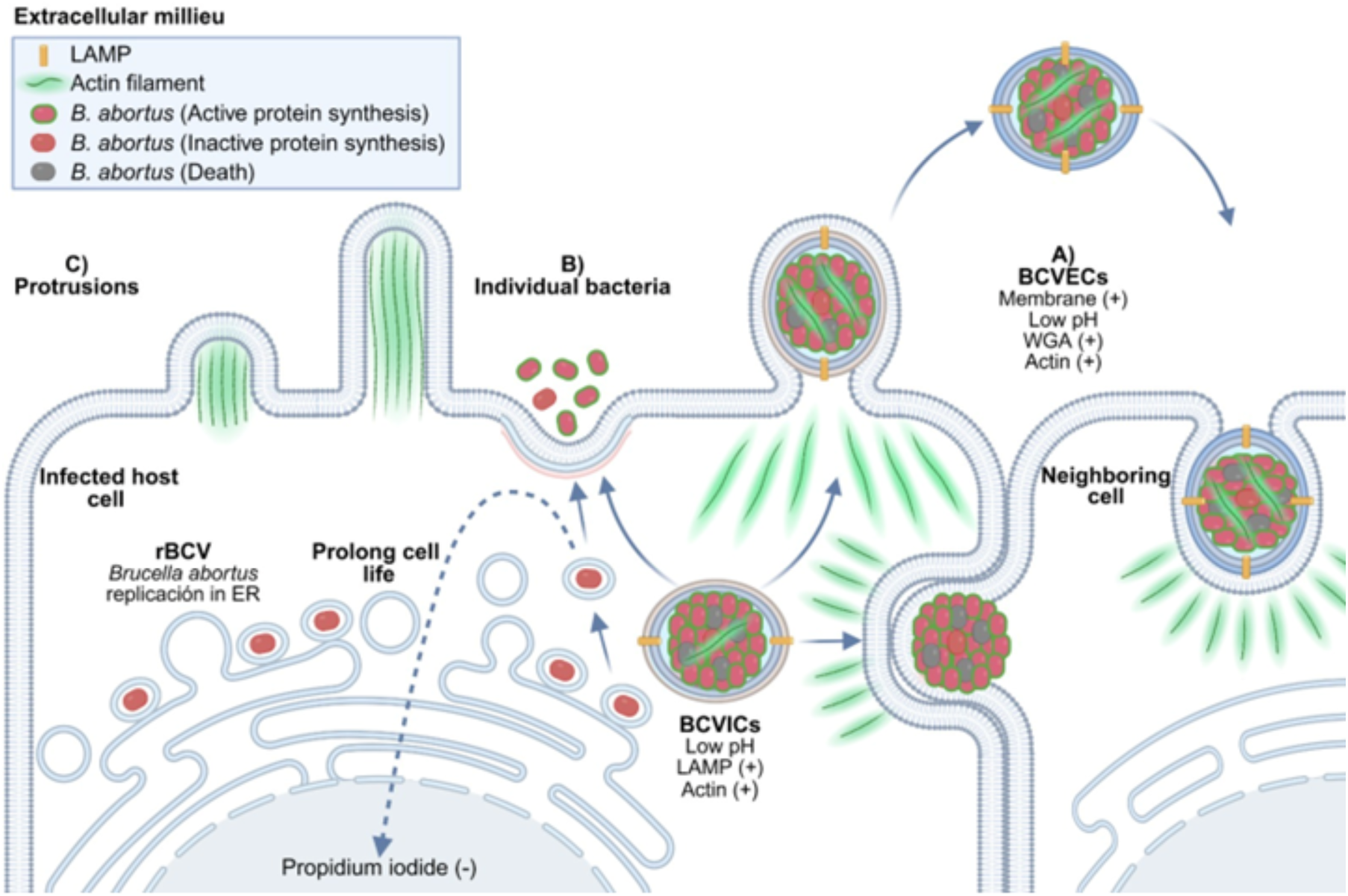
Model illustrating the dynamics of *B. abortus* release from previously infected cells. Following extensive replication in endoplasmic reticulum-derived compartments, *B. abortus* organisms assemble into an acidic vacuole with autophagosomal characteristics (BCVICs), as proposed by Starr et al. (2012). During this period, replicating bacteria prolong the lifespan of the infected cell. (A) Dissemination through BCVECs. The BCVEC formation is initiated by the activation of Rho subfamily GTPases and the active polymerization of actin filaments. (B) Dissemination by individual bacteria. Free *B. abortus* organisms are released into the extracellular milieu. These single bacteria can reach adjacent cells or distant sites. (C) Protrusion phenomenon.

We do not know why some infected cells primarily generate BCVIC while others display dispersed bacteria throughout the cytoplasm, not only observed in cultured HeLa cells but also other epithelial cells, such as placental trophoblasts (5,18,14). Certainly, the tendency to form large BCVIC increases over time, possibly related to the crowded effect within the cytoplasm. It is known that crowding can affect not only diffusion rates and macromolecular interactions but also cell volume regulation, all of which impact intracellular vacuole dynamics (27). Once the BCVIC reaches a given size, ranging from 15 to 650 μm^3^, the cell is compelled to release the bacterial clumps. However, larger BCVEC above 250 μm^3^ are rare, suggesting that the process of shedding smaller BCVEC (e.g., <100 μm^3^) is constitutively sustained.

The close packing of bacteria within BCVEC suggest that the *Brucella* coccobacilli follow the rules of the closest-packed configuration, which depends on the number and bacteria size (16). At least for less than 50 brucellae coccobacilli, the packing structures are arranged in polyhedrons of various shapes. For instance, the packing of 12 coccobacilli approximates a cube (16). In contrast, the packing of 50 and 100 coccobacilli approximates spherical configurations. Nevertheless, the packing structures significantly differ. Although larger BCVEC may maintain spherical shape, their bacterial arrangements will vary depending on the vacuole size (16). Higher numbers of bacteria are predicted to make the BCVEC less stable and prone to disruption. Indeed, the BCVIC and BCVEC, with more than 250 bacteria, are less abundant than those with fewer bacteria.

Despite being ordered within BCVEC, there is bacterial heterogeneity regarding dead and live organisms, as well as the metabolic activity of the former. Although most live bacteria inside BCVEC remain metabolically active, a small proportion are inactive, and nearly 20-25% are dead bacteria. While metabolically live bacteria enhance their ability to infect other cells, dead bacteria are likely donors of cohesive material such as DNA and histone-like proteins (e.g., MucR), potentially acting as “glue” for bacterial aggregation (28,29). This tight clumping probably forms microcolonies that shield against bactericidal agents and facilitate quorum sensing for initiating a new infection cycle (30).

From the standpoint of composition (LAMP-1, actin filaments, acidity and size), BCVIC and BCVEC display very similar properties, and it is well known that intracellular acidity triggers the expression of genes necessary for the bacteria’s survival and replication inside host cells (31). Indeed, *Brucella* BCVIC autophagic characteristics may be related to mechanisms that activate virulent processes, enhancing bacterial infectivity and adhesion as described previously (31). Moreover, bacteria isolated directly from infected cells at later stages exhibit enhanced virulence, as demonstrated by higher adherence and penetration capabilities, compared to those grown in bacteriological cultures (31). This phenomenon is linked to the BvrR/BvrS system, which recruits VjbR and VirB virulence factors (31).

It is not known whether brucellae within the BCVIC promote their exit through type IV bacterial effectors that eventually trigger cytoskeletal polymerization through the action of small GTPases. It was observed that heavily infected cells produce cytoplasmic protrusions accompanied by the polymerization of cytoskeletal actin proteins, resulting from the persistent local recruitment of small GTPases of the Rho subfamily. The fact that LAMP-1 and actin remain closely associated with BCVEC indicates the ubiquity of this process during the *B. abortus* intracellular biogenesis. The actin-rich filopodia-like protrusions may also facilitate the intercellular bacterial infection (32), a phenomenon that we have concurrently observed in *B. abortus* infected cells and documented in an accompanying publication (33). Analog to the initial local recruitment of small GTPases for the entrance of invading *Brucella* organisms to HeLa cells (4), the exit o*f B. abortus* is also promoted by the recruitment of GTPases of the Rho subfamily. The recruitment of Rho GTPases, which control the dynamics of the cytoskeleton, could be a ubiquitous mechanism for the release of large cell aggregates, as they are also required for the protrusion of apoptotic bodies (34), a phenomenon reminiscent of the release of BCVEC.

It has been established that vacuolar lytic enzymes intracellularly degrade autophagic bodies, and the degradation products are reused or released (35,36). However, this is not true for the BCVIC, which exhibits autophagic characteristics (8). In *Chlamydia trachomatis* intracellular parasitism, membrane-bound bacterial clusters budding from the cells are released through the concourse of GTPases of the Rho subfamily and the concomitant recruitment of actin filaments (20). As in *Chlamydia*, the generation of cytoskeletal filaments in *Brucella*-infected cells, associated with the formation of cytoplasmic extensions, is also linked with the BCVEC protrusion. However, chlamydial infection vesicles lack autophagosome characteristics, are not acidic, and do not contain LAMP-1; instead, they display properties of early endosomes (37).

Intracellular *B. abortus* organisms within neutrophils also promote their infectivity towards macrophages that ingest these infected polymorphonuclear cells. Once inside macrophages, the *Brucella*e organisms replicate significantly more than when macrophages are infected with bacteria alone (38,39). Similar to *Brucella*-infected neutrophils, the BCVEC are internalized by other cells and initiate a new infection cycle with a substantial bacterial load. As in neutrophils, these bacteria also seem highly virulent (38, 31).

The ability of a proportion of *B. abortus* to exit encased within a host-cell-derived membrane through mechanisms that do not involve cell death provides significant advantages. Indeed, the acidic membrane-bound BCVEC with the characteristics of autophagosomes or apoptotic bodies may be internalized by professional phagocytes, such as macrophages, neutrophils, or dendritic cells, thereby suppressing proinflammatory signals (40–42). In contrast to naked bacteria readily recognized by the immune system, membrane-bound BCVEC may evade immune detection. It remains to be solved whether the membrane-bound BCVEC are more infectious than the naked ones or vice versa. It is plausible that the naked BCVEC are more active in infecting neighboring cells, while the membrane-bound BCVEC travel larger distances, evading antibodies, complement, or other bactericidal substances in the extracellular milieu. These phenomena would enable *Brucella* organisms to maintain their stealthy strategy, establish a persistent infection, and facilitate dissemination to other cells and hosts (15).

## Materials and Methods

### Bacterial strains and culture

*B. abortus* 2308, expressing a constitutive red fluorescent protein (RFP) from Discosoma coral (*B. abortus*-RFP) (Unité de Recherche en Biologie Moléculaire, Facultés Universitaires Notre-Dame de la Paix, Namur, Belgium), *B. abortus* 2308W expressing a constitutive green fluorescent protein (*B. abortus*-GFP) (42,43) and *B. abortus* 2308W with the anhydrotetracycline (ATc)-inducible pJC45-gfp plasmid (*B. abortus* 2308W-pJC45-gfp) were used in this study (44). In addition, *B. abortus*-RFP was electroporated with the anhydrotetracycline (ATc)-inducible pJC45-gfp plasmid (*B. abortus*-RFP-pJC45-gfp) following previous protocols (44). *Brucella* strains were grown on trypticase soy agar (TSA), inoculated in 20 mL of Trypticase Soy Broth, and incubated at 37 °C with constant shaking for 24 h. Colony-forming units per mL (CFU/mL) were estimated by the absorbance of the culture in TSB at 420 nm and measured using a previously standardized formula (31). All the *B. abortus* strains used displayed a similar replicating pattern in cells and were fully virulent.

### Infection of cell cultures

HeLa ATCC clone CCL-2 cells were cultured and infected with *B. abortus* 2308W-pJC45-gfp*, B. abortus*-RFP, *B. abortus*-RFP-pJC45-gfp or *B. abortus*-GFP as described previously (31). Briefly, for fluorescence analysis, HeLa cells were seeded on 13-mm glass slides in 24-well tissue culture plates two days before infection to obtain a final concentration of 5 × 10^5^ cells per well. For live imaging, 96-well HeLa cells were seeded two days before infection to obtain a final concentration of 3 × 10^4^ cells per well previous to analysis using a BioTek Lionheart FX Automated Microscope (Agilent). A multiplicity of infection (MOI) of 500 was determined by estimating the bacterial population using optical density, followed by diluting the bacteria in Modified Eagle’s Medium (DMEM) supplemented with 5% fetal bovine serum. Infected cells were centrifuged at 330 *× g* for 5 min at 4 °C, incubated at 37 °C and 5% CO2 for 45 minutes. Each well was washed in triplicate with 1× PBS and incubated for 1 hour with DMEM medium supplemented with 100 μg/mL gentamicin to remove extracellular bacteria. The medium was then replaced with DMEM maintenance medium supplemented with 5 μg/mL gentamicin and incubated for the indicated hours in each assay.

### Intracellular and extracellular bacterial quantitation

Quantitation of intracellular and extracellular *B. abortus* organisms was performed as described before ^31^. Briefly, the DMEM maintenance medium supplemented with 5 μg/mL gentamicin in the cell culture was discarded and replaced with DMEM medium without gentamicin. After incubation for 4 hours, the cell supernatants were taken, and serial dilutions were performed and plated in Tryptic Soy Agar. Concurrently, intracellular bacteria were determined at the same intervals. The infected cells were disrupted with a 0.1% Triton X-100 solution for 10 minutes. Quantitation was performed either by direct counting of red fluorescent bacteria under the microscope or through dilutions followed by plating on TSA and culturing for 48 hours. This procedure was repeated 6, 29, 48, 54, 70, and 100 hours post-infection (p.i.).

### Purification of BCVIC and BCVEC

HeLa cells seeded in a 6-well plate were infected, and at 48 hours p.i., the cell culture maintenance medium was changed to a medium without gentamicin to allow BCVEC to exit from infected cells. The supernatant with BCVEC and free bacteria was then collected. Thereafter, the infected cells were washed with DMEM, lysed with 0.1% Triton X-100 in 1 × PBS, and the suspension was collected after 10 minutes of incubation. Supernatants and cell lysates were used to purify BCVIC and BCVEC, using the same protocol. The suspensions were centrifuged at room temperature at 90 *x g* for 5 minutes. The resulting supernatants were centrifuged at 300 *x g* for 5 minutes at room temperature. The sediment obtained from the second centrifugation, containing purified BCVIC or BCVEC, was processed for analysis using fluorescence or electron microscopy, as described below.

### Fluorescence microscopy

Fluorescence microscopy was performed as described previously (4,31). *B. abortus*-infected HeLa cells or purified vesicles were fixed with ice-cold 3% paraformaldehyde for 10 minutes. Then, cells and vesicles were washed in 1 × PBS and incubated for 10 min with 1 × PBS containing 50 mM NH_4_Cl. To stain DNA in cell nuclei and bacteria, infected cells were incubated for 10 minutes with 1 × PBS containing 0.5 µg/ml diamidino-2-phenylindole (DAPI) (Thermo Fisher Scientific). Samples were processed with different treatments depending on the experiment (as indicated in the figures). Cells or vacuoles were stained with 6.6 µM Alexa Fluor 488-phalloidin (Invitrogen™) in a 10% horse serum solution for 30 minutes to detect actin. BCVIC and BCVEC were stained with a 0.5 µg/mL germ-wheat agglutinin conjugated with Alexa Fluor 488 (GWA-AlexaF-488, Invitrogen™). BCVIC and BCVEC were labeled with a 1 µg/mL LAMP1 rabbit monoclonal antibody H4A3 (Abcam) for 30 minutes and 2 µg/mL Alexa Fluor 488 anti-rabbit conjugate (Invitrogen™) for 30 minutes. To determine the acidity of the protruding BCVEC, infected HeLa cells were incubated with 50 nM LysoTracker Red DND-99 (Invitrogen™) at 48 hours post-infection for 1 hour before analysis.

We followed a previously described procedure to determine if BCVIC and BCVEC are enclosed within an impermeable host membrane (31). Briefly, to label bacteria found on membraneless vesicles, BCVEC and BCVIC cells were washed three times with 1× PBS and incubated for 30 min at 4°C with an in-house cow IgG conjugated with fluorescein isothiocyanate (FITC), mostly reacting against the *Brucella* smooth LPS, as evaluated by Western blot (13). Cells were then fixed with ice-cold 3% paraformaldehyde for 10 min. Samples were washed once, incubated for 10 min with 1× PBS containing 50 mM NH4Cl, and permeabilized for 10 min with 0.1% Triton X-100. After incubation, slides were washed three times with 0.1% Triton X-100, and the nuclei were stained with 0.5 μg/ml DAPI (Thermo Fisher Scientific). Finally, slides were mounted using ProLong gold antifade mountant (Invitrogen**™**) before analysis by fluorescence microscopy. Before and after each experiment, cells were examined with 1 µg/mL Hoechst (Abcam) to preclude *Mycoplasma* contamination.

For live imaging analysis, infected cells, BCVIC and BCVEC were recorded using a BioTek Lionheart FX Automated Microscope for up to 100 hours. Confocal imaging was performed with a Nikon A1R HD25 laser-scanning microscope equipped with ×60 (Plan Apo λ Oil) and ×100 (SR HP Apo TIRF) objectives. The microscope was equipped with a Nikon A1 LFOV camera and four laser lines 405 nm (20mW), 488 nm (70 mW), 561 nm (70 mW), and 647 nm (125 mW) and the following emission/excitation wavelengths: DAPI 450/405 nm; FITC 525/488 nm; TRICT 595/561 nm. Images were acquired with NIS-Elements advanced workstation software 5.42.04 (Nikon). All images were captured in 12-bit format with a resolution of 1024 x 1024 pixels at room temperature. When stated, raw images were processed using the Denoise.ai module.

### Size determination of intra- and extracellular clumps

The purified, fixed, and stained BCVIC and BCVEC were observed under a confocal fluorescence microscope using the above-described software. The diameters of the vesicles were measured, and their volume was estimated. Likewise, the average sizes and volume of *B. abortus* were determined by measuring bacteria in photographs taken through transmission and scanning microscopy. Based on these measurements, the average volume of *B. abortus* coccobacilli was estimated to be 0.49 ± 0.04 μm^3^. The number of bacteria in each vesicle was estimated using the formula: vesicle volume µm³/*B. abortus* average volume µm³ × 0.74 (the estimated proportion of space filled by roads packed within a sphere). For the estimation of the packing order of bacteria within BCVEC, photographs of the vesicles were amplified, and the number of fluorescent spots was counted using the Adobe Photoshop ® density color pixel function of the image’s sections and the method for the close packing of rods on a spherical surface followed (16).

### Determination of live, dead, and metabolically active *B. abortus*

*B. abortus*-GFP, *B. abortus* 2308W-pJC45-*gfp*, and *B. abortus*-RFP-pJC45-*gfp* were used to evaluate live and dead bacteria and protein synthesis in “metabolically” active bacteria. After the purification of BCVIC and BCVEC, GFP expression of *B. abortus*-pJC45*-gfp* or *B. abortus*-RFP-pJC45-*gfp* was induced by adding 250 nM B-anhydro-tetracycline for 6 hours. *B. abortus*-RFP-pJC45-*gfp* displaying red and green fluorescence were considered alive and metabolically active in protein synthesis. Likewise, *B. abortus* 2308W-pJC45-*gfp* displaying green fluorescence were considered metabolically active in protein synthesis. *B. abortus*-RFP-pJC45-*gfp* displaying only red fluorescence was considered alive but metabolically inactivated. Bacteria stained with 1X PBS containing 1µg/mL propidium iodide were recorded as dead bacteria ^17^. All bacteria (dead and alive) stained blue with 1X PBS containing 0.5 ug/ml DAPI. For some experiments, the purified BCVIC and BCVEC were disrupted with 0.1% Triton X-100 dissolved in 1X PBS before counting the bacteria under the fluorescent microscope. The ratio of red fluorescence versus green fluorescence per bacteria was estimated by using the Histogram function of Adobe Photoshop program of 50 single *B. abortus*-RFP-pJC45-*gfp* expressing or not GFP protein (17).

### Analysis of cells, BCVIC and BCVEC by electron microscopy

The pellets obtained from purified BCVEC and HeLa cells infected were fixed with a modified Karnovsky solution (2% glutaraldehyde, 2% paraformaldehyde in 0.05 M phosphate buffer, pH 7.2, for two hours) and then washed with phosphate buffer. Then, they were post-fixed with 1% osmium tetroxide fixative and washed with distilled water. Subsequently, the sample was dehydrated using ethanol gradients (30%, 50%, 70%, 80%, 90%, 95%, and 100%) for 15 minutes. Samples were dried, and finally, the slides with infected cells were mounted on an aluminum base and recovered with gold (12 nm) in a Sputter coater Denton Vacuum model DESK V (Denton Vacuum, Inc.). Cells were analyzed using an electron scanner JEOL JSM-6390 LV with an acceleration voltage between 10 and 20 kV, and images were taken at different magnifications.

Transmission electron microscopy was performed as described previously, with slight modifications (4). Briefly, *B. abortus* 2308W-infected HeLa monolayers were fixed with 2.5% glutaraldehyde and 2% paraformaldehyde in 0.1 M phosphate buffer. Fixed cells were incubated at 40°C for 5 minutes, and 100 μL of 3% low-melting agarose at 40°C was added. The temperature was lowered, and five volumes of 2.5% glutaraldehyde in PBS were added to the solid agarose block and incubated overnight at 4°C. Samples were placed in 1% OsO_4_ solution for one hour post-fixation, dehydrated in a graded ethanol concentration, and infiltrated with Spurr resin. Thin sections on 300 mesh collodion-coated grids were stained with uranyl acetate and lead Sato’s solution. Preparations were examined with a Hitachi H-7100 electron microscope operating at 75 kV.

### Participation of small GTPases of the Rho subfamily

Three ng/ml cytotoxic necrotizing factor 1 (CNF) or 1 µg/ml cytochalasin was added to infected cells at 48 hours p.i., for 2 hours. Cells were washed with 1 × PBS at 72 hours p.i., fixed with paraformaldehyde or modified Karnovsky solution, and prepared for fluorescence or electron microscopy analysis as previously described. Wells without CNF or cytochalasin were used as controls. Additionally, at 72 hours p.i., the DMEM maintenance medium, supplemented with 5 μg/mL of gentamicin, was changed to a gentamicin-free medium to promote bacterial egress from infected cells. After 4 hours of incubation, supernatants from infected cells were transferred, serially diluted, and plated in TSA. After washing the infected cells with 1X PBS, the cells were lysed with 0.1% Triton X-100 for 10 minutes. The supernatant was then diluted, and bacteria were counted by plating them in TSA and culturing for 48 hours at 37 °C, as described previously (31).

For live imaging analysis, 3 ng/ml cytotoxic necrotizing factor 1 (CNF) or 50 ng/ml *Clostridioides difficile* toxin B (TcdB) was added to infected cells at 48 hours p.i., for 2 or 1 hour, respectively. Cells were washed with 1 × PBS and BCVIC and BCVEC were recorded using a BioTek Lionheart FX Automated Microscope for up to 100 hours as described above. Wells without CNF or TcdB were used as controls.

### BCVEC infection assessment

The maintenance medium with gentamicin from infected HeLa cells was changed to DMEM medium without gentamicin or maintained with gentamicin at 48 hours p.i. The proportion of newly infected cells through released BCVEC was assessed through live imaging using a BioTek Lion Heart Imager. Cell nuclei were stained with DAPI as described above.

### Statistical Analysis

Student t-test, the Kruskal-Wallis, and Dunn’s multiple-comparison tests were used accordingly. The significance was indicated as: *, *p*< 0.05; **, *p* < 0.005.

